# Learning maximally spanning representations improves protein function annotation

**DOI:** 10.1101/2025.02.13.638156

**Authors:** Jiaqi Luo, Yunan Luo

## Abstract

Automated protein function annotation is a fundamental problem in computational biology, crucial for understanding the functional roles of proteins in biological processes, with broad implications in medicine and biotechnology. A persistent challenge in this problem is the imbalanced, long-tail distribution of available function annotations: a small set of well-studied function classes account for most annotated proteins, while many other classes have few annotated proteins, often due to investigative bias, experimental limitations, or intrinsic biases in protein evolution. As a result, existing machine learning models for protein function prediction tend to only optimize the prediction accuracy for well-studied function classes overrepresented in the training data, leading to poor accuracy for understudied functions. In this work, we develop MSRep, a novel deep learning-based protein function annotation framework designed to address this imbalance issue and improve annotation accuracy. MSRep is inspired by an intriguing phenomenon, called neural collapse (NC), commonly observed in high-accuracy deep neural networks used for classification tasks, where hidden representations in the final layer collapse to class-specific mean embeddings, while maintaining maximal inter-class separation. Given that NC consistently emerges across diverse architectures and tasks for high-accuracy models, we hypothesize that inducing NC structure in models trained on imbalanced data can enhance both prediction accuracy and generalizability. To achieve this, MSRep refines a pre-trained protein language model to produce NC-like representations by optimizing an NC-inspired loss function, which ensures that minority functions are equally represented in the embedding space as majority functions, in contrast to conventional classification methods whose embedding spaces are dominated by overrepresented classes. In evaluations across four protein function annotation tasks on the prediction of Enzyme Commission numbers, Gene3D codes, Pfam families, and Gene Ontology terms, MSRep demonstrates superior predictive performance for both well- and underrepresented classes, outperforming several state-of-the-art annotation tools. We anticipate that MSRep will enhance the annotation of understudied functions and novel, uncharacterized proteins, advancing future protein function studies and accelerating the discovery of new functional proteins. The source code of MSRep is available at https://github.com/luo-group/MSRep.

## 1 Introduction

The advancement of next-generation sequencing has led to an unprecedented accumulation of genomic data, far outpacing our capability to elucidate the exact function of proteins encoded in sequenced genomes. While protein function annotation is a central task in life sciences, the level of scientific investigation varies greatly between proteins. Some, like p53, have been the focus of extensive study for decades, while many others remain largely understudied,^1, 2^ a disparity gap that hinders progress in biomedical research.^3^ As of 2023, nearly 80% of the 248 million proteins in the UniProt database^4^ remain without precise function annotation.^5^ Moreover, within the human proteome, roughly 95% of life science publications today focus on just a subset of 5,000 well-studied proteins like p53.^6^ Even in a minimal synthetic organism *Mycoplasma* JCVI-3 with *<*400 genes, the functions of 30% of encoded proteins remain unknown.^7^ Additionally, while ~ 3,000 human proteins are expected to be potential drug targets, only 10% are currently targeted by FDA-approved drugs.^8^ More importantly, this growing annotation imbalance creates a ‘rich-get-richer’ problem since new research preferentially targets proteins that are already well-studied,^9^ driven by reasons such as experimental feasibility,^3, 10^ availability of prior knowledge,^2^ and scientists’ perceptual bias.^10^ Recognizing these challenges, initiatives such as the Understudied Proteins Initiative have been established, advocating for the systematic study and characterization of understudied proteins.^11, 12^

Computational approaches have emerged as a cost-effective solution to support experimental techniques to annotate functions for understudied proteins. Established algorithms include BLASTp^13^ and profile hidden Markov models (pHMMs)^14^ leverage sequence alignment to transfer annotations from annotated to uncharacterized proteins. Recent rapid advances in machine learning (ML), particularly deep learning, have significantly improved protein function prediction by learning from patterns across protein families and directly predicting functions from sequences or structures. ML approaches have shown remarkable success across various tasks, including the predictions of Gene Ontology (GO) terms,^5, 15–24^ Enzyme Commission (EC) numbers,^5, 21, 23, 25–30^ Pfam labels,^19, 21, 31, 32^ and in the Critical Assessment of Functional Annotation (CAFA).^33, 34^ The impacts of ML methods by their integration into major biological databases such as UniProt^4^ and Pfam,^35^ where ML-predicted functions are now part of official function annotation releases.^36, 37^

Despite these advances, ML methods for protein function prediction still perform poorly for rare functions (e.g., those associated with *<*10 proteins) or proteins lacking homologs.^38^ The root of this problem lies in the imbalanced distribution of function annotations within existing databases like GO terms,^39^ EC numbers,^40^ Pfam labels,^35^ and more. Due to annotation bias, these databases often exhibit a long-tail distribution (Fig 1a), where a few function classes are associated with the majority of proteins (‘head classes’), while many others are sparsely annotated (‘tail classes’). Accurate predictions for tail classes are especially valuable, as they often correspond to novel or understudied proteins. However, ML models trained to minimize the overall loss on imbalanced data tend to allocate most of their “learning capacity” to overly focus more on optimizing prediction accuracy for head-class proteins than tail classes. Paradoxically, while the goal of ML-based function prediction tools is to reduce annotation bias, this very bias becomes the key obstacle, preventing ML models from realizing their full potential and leading to significantly lower prediction accuracy for understudied proteins compared to well-characterized ones. Some studies have attempted to address this imbalance, such as using inverse class frequency or prediction confidence to up-weight understudied proteins and down-weight already well-classified proteins in model training,^20, 41^ but these methods remain highly sensitive to the large variance in class sizes. Another line of work, including ours, has employed contrastive learning and one-vs-all nearest neighbor search, implicitly giving equal weights to all classes.^23, 28, 42^ While promising for rare classes, these methods are still vulnerable to bias from the frequent sampling of head-class proteins during contrastive learning, skewing the learned representation space toward well-studied proteins.

**Figure 1:**
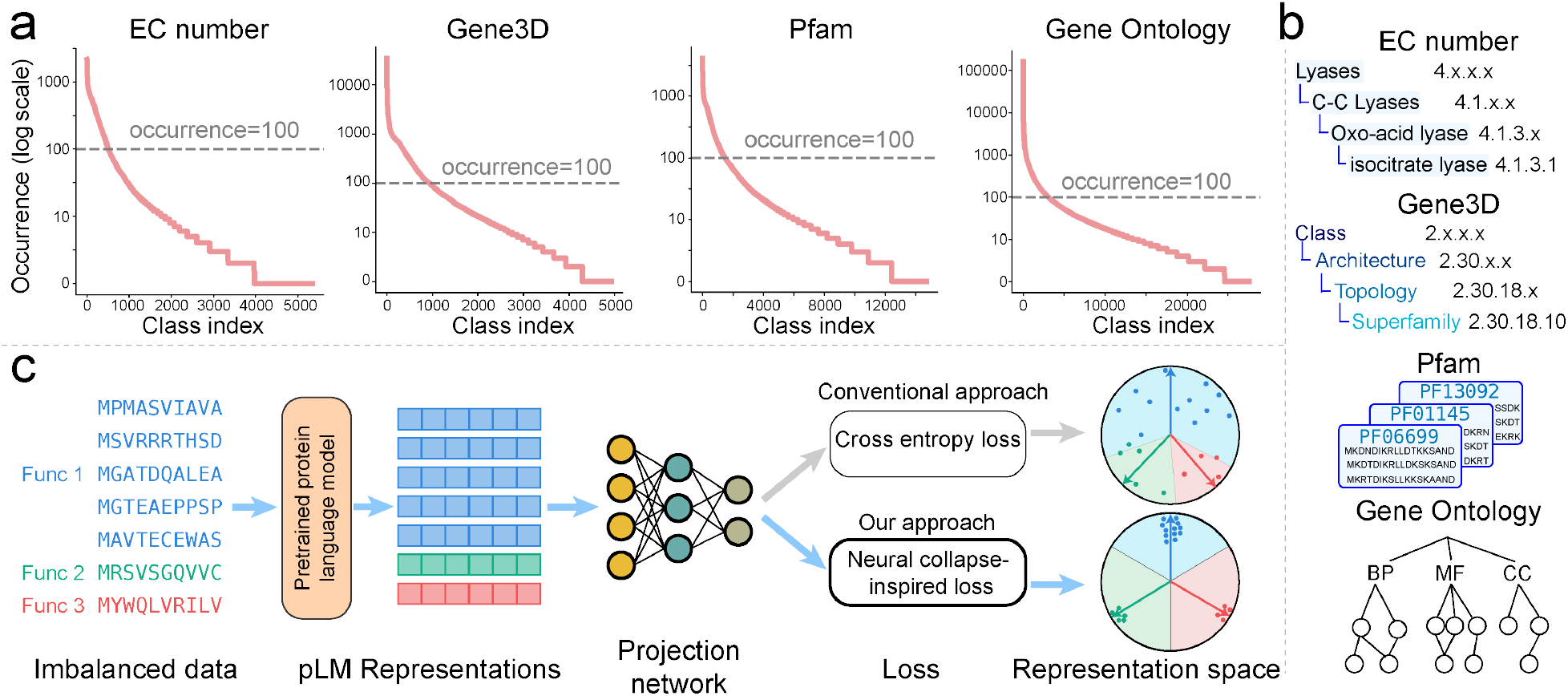
Overview of MSRep. (**a**) Long-tail distribution of function class occurrences in four protein function annotation schemes, including EC numbers, Gene3D codes, Pfam families, and Gene Ontology terms. The occurrence of a function class is defined as the number of proteins associated with it. Statistics were calculated based on function annotations in UniProt/Swiss-Prot^51^ as of 2022/05/25. (**b**) Illustrations of the four protein function annotation schemes. BP: biological process, MF: molecular function, CC: cellular component. (**c**) The schematic overview of MSRep in comparison to the conventional classification approach (grey arrows). In the representation space, the blue, green, and red arrows represent the class centers of different function classes, with colored dots indicating the embeddings of proteins with these functions. The shaded regions correspond to the parts of the representation space classified to each function class. Func: function, pLM: protein language model.

To address these challenges, we develop MSRep (maximally spanning representation learning), a deep learning method that accounts for protein annotation bias and delivers accurate, unbiased protein function predictions. Recognizing that existing ML methods often prioritize accuracy for overrepresented function classes at the expense of rare ones, MSRep aims to correct this imbalance by allocating equal “parameter budgets” to model all function classes. MSRep is inspired by the intriguing phenomenon of *Neural Collapse* (NC), a consistent characteristic observed in high-performance deep learning classification models.^43, 44^ NC states that, in the last layer of a well-trained neural network, embeddings of samples from the same class collapse to their class-specific means, while embeddings from different classes are maximally separated. Since NC is a hallmark in high-accuracy models across various classification tasks with balanced data,^43^ we hypothesize that explicitly inducing NC in models trained on imbalanced data, such as protein function annotations, can significantly enhance predictive accuracy for minority classes. To achieve this, MSRep refines a pre-trained protein language model (pLM)^45^ to exhibit NC-like behavior, where protein embeddings annotated with the same function collapse together, while maintaining equal, maximal separation from other function classes. This ensures that tail-class proteins occupy an embedding space equal in size to that of head classes, preventing the model from overfitting to overrepresented classes or proteins during training.

We thoroughly evaluate MSRep across four protein function annotation tasks: EC number, structural domain, protein family, and GO term predictions. In all tasks, MSRep outperforms state-of-the-art protein function predictors, significantly improving annotation accuracy for both well-studied and understudied functions. MSRep’s representation learning framework can be readily adapted to other protein function annotation tasks that can be formulated as multi-label classification problems. We anticipate that MSRep will contribute to the systematic efforts of reducing the annotation gap by associating uncharacterized proteins with proteins of known functions,^11, 12^ empowering the discovery and engineering of functional proteins.

## 2 Methods

### 2.1 Problem setup

Multiple protein function classification schemes (Fig. 1b) have been developed to annotate proteins’ functions, such as the Gene Ontology^46^ (general biological functions), EC numbers^40^ (enzymatic activities), Pfam^35^ (protein family), Gene3D^47^ (structure domain), along with other databases describing protein interactions^48^ and pathways.^49^ The goal of *protein function annotation* is to assign one or more labels from a controlled vocabulary of functions (e.g., GO terms or EC numbers) that describe a protein’s functional roles. In this work, we consider the sequence-based protein functions prediction, where the input is a protein’s amino acid (AA) sequence, though the setting can be readily extended to structure-based function prediction using 3D protein structures as input.^20^

In the context of ML, protein function annotation is a multi-class, multi-label classification problem. Denote *𝒞* = {1, 2, …, *K*} as the predefined vocabulary of function labels in a function annotation scheme (e.g., GO terms or EC numbers), and ***s*** = *a*_1_*a*_2_ … *a*_*L*_ as a protein sequence of length *L*, where *a*_*i*_ ∈ *𝒮* is the amino acid at position *i*, and *𝒮* is the set of 20 canonical AAs. A protein function annotation database can be represented as a list of protein-annotation pairs 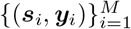 where ***y***_*i*_ is the set of function annotations for sequence ***s***_*i*_. Since a protein may perform multiple functions, ***y***_*i*_ may contain multiple labels from *𝒮*.

Our goal is to learn a predictive ML model *f*_*θ*_ : *𝒮* → 2^*C*^ that predicts one or more function labels for a protein based on its sequence. Existing leading ML methods for protein function prediction are typically deep learning models.^21, 23, 28–31^ These models generally decouple into two modules *f*_*θ*_ = *g*_*ω*_ ∘ *h*_*ϕ*_, including a feature extractor *h*_*ϕ*_, which maps the input sequence to an embedding ***x*** = *h*_*ϕ*_(***s***) ∈ ℝ^*d*^, and a linear classifier *g*_*ω*_, which predicts the likelihood of the protein belonging to each of the *K* function classes, ***ŷ*** = *g*_*ω*_(***x***) ∈ ℝ^*K*^.

Interestingly, several recent models omit the explicit learning of the classifier *g*_*ω*_. Instead, they rely on the protein proximity induced by the embedding space learned by the feature extractor *h*_*ϕ*_ to infer functions, following the “guilt-by-association” principle, i.e., similar proteins are likely to share similar functions.^23, 28, 50^ For example, CLEAN,^28^ a state-of-the-art enzyme function predictor we previously developed, employs contrastive learning to train *h*_*ϕ*_, generating a sequence embedding space where proteins with similar functions are pulled together, while those with different functions are pushed apart. Function prediction is then performed via nearest neighbor search in this space, transferring function labels from annotated proteins to the query protein, without explicitly training a classifier *g*_*ω*_. This strategy has been extended by the Protein-Vec method to other tasks, such as GO term and structure domain classification.^23^

While contrastive learning has proven effective in these studies, it only optimizes local pairwise embedding distances between proteins, without capturing group-wise relationships that reflect similarity across function classes. Furthermore, it does not address the class imbalance issue in protein function annotations. Our approach differs conceptually, as we aim to optimize the global distribution of protein embeddings, ensuring a balanced allocation of the embedding space to mitigate the function class imbalance.

### 2.2 MSRep: Maximally spanning representation learning for protein function annotation

We propose MSRep, a neural network that learns maximally spanning representations of proteins for function annotations (Fig. 1c). MSRep specifically accounts for the bias in protein annotation data during representation learning, enhancing performance across various protein function prediction tasks. Receiving a protein’s sequence as input, MSRep uses ESM-1b,^45^ a pre-trained protein language model (pLM), to generate an initial embedding for the protein. Although pLM embeddings have proven to be information-rich and effective in many protein biology tasks, they do not explicitly address the class imbalance issue in protein function annotation. To tackle this, MSRep refines ESM embeddings through a multi-layer perceptron (MLP) “adapter” network, which reprojects the sequence embeddings into a new representation space that satisfies the NC properties. In this space, embeddings of proteins with the same function cluster around their class mean, while the mean embeddings of all classes are maximally separated, forming equal-sized angles between any given pair (Fig. 1c). During training, only the MLP parameters are optimized, while ESM remains frozen. This strategy is computationally efficient, leveraging the strong priors important to function captured by the pre-trained ESM–such as sequence, structural, and evolutionary features–while avoiding the need to retrain the entire model from scratch. At the prediction (inference) stage, given a query protein, MSRep performs a nearest-neighbor search against proteins with known functions in the training data and transfers the function annotations from the protein with the closest embedding distance to the query. Below we review the key concepts of NC and describe how we leverage NC to develop MSRep.

#### 2.2.1 Neural Collapse: Revisit

The recent work by Papyana et al.^43^ made a remarkable observation with compelling mathematical insights. It states that well-trained neural networks with the cross-entropy loss on balanced classification tasks typically exhibit the Neural Collapse (NC) phenomenon in the last layer, where the learned embeddings of samples from the same class collapse to their within-class mean embeddings (i.e., class centers). Meanwhile, these class centers, when zero-centered, are maximally separated, with equal-sized angles between any pairs in the high-dimensional embedding space. Equivalently, this structure can be described as a simplex equiangular tight frame (ETF),^52^ formally defined as follows.

##### Definition 1

(Simplex Equiangular Tight Frame). A set of vectors 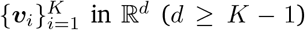 is called a simplex equiangular tight frame (ETF) if ∥***v***_1_∥ = ∥***v***_2_∥ = · · · = ∥***v***_*K*_∥ and

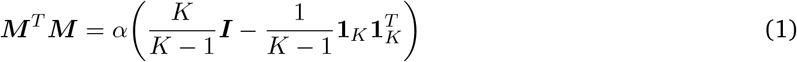

where ***M*** = [***v***_1_, ***v***_2_, · · ·, ***v***_*K*_] ∈ ℝ^*d×K*^, *α* is a non-zero constant, ***I*** ∈ ℝ^*K×K*^ is the identity matrix, and **1**_*K*_ ∈ ℝ^*K*^ is the all-one vector. Intuitively, a simplex ETF is a set of vectors of equal lengths, maximally separated from each other by equal angles.

Formally, consider a training set of *n* data points and let the class set be *𝒞* = {1, 2, …, *K*}. Denote the last-layer embeddings as ***x***_1_, ***x***_2_, …, ***x***_*n*_ ∈ ℝ^*m*^, the arithmetic mean of the embeddings for each class as ***µ***_1_, ***µ***_2_, …, ***µ***_*K*_, and the final linear classifier of the neural network as *g*_*ω*_(***x***). For simplicity, we define a helper function *ψ* : ℝ^*m*^→ *𝒞*, which maps the embedding ***x*** to its ground truth class label *ψ*(***x***) = *c* ∈ *𝒞*. Here, we consider the case where each sample ***x*** has only one class label, but this will later be extended to the multi-label setting required for protein function annotation, where one sample can have multiple labels. The NC phenomenon can be formally described by the following four properties:

##### (*NC*_1_) Variability collapse

In the terminal phase of training, the intra-class embedding variances collapse to zero, and the embeddings of samples within the same class collapse to their class mean, i.e.,

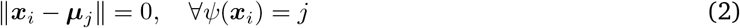

##### (*NC*_2_) Convergence to simplex ETF

The class means of all the *K* classes, after zero-centering normalization, converge to form an ETF (Definition 1). Let 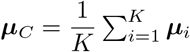, then we have

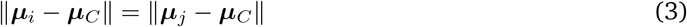

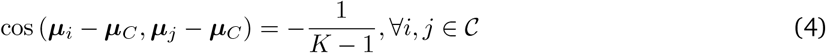

Geometrically, the class means converge to having equal length and being maximally separated from each other, with equal-sized angles between any given pair.

##### (*NC*_3_) Convergence to self-duality

The weights of the linear classifier *g*_*ω*_(***x***) become parallel to the corresponding zero-centered class means, up to a constant factor rescaling. Let ***w***_*j*_ denote the *j*-th row of the weights matrix *g*_*ω*_ as ***w*** ∈ ℝ^*K×d*^, then:

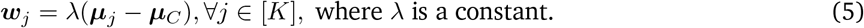

##### (*NC*_4_) Simple classification rule

For a new data point with last-layer embedding ***x***^*′*^, the prediction made by the classifier is equivalent to selecting the class center ***µ***_*j*_ closest to ***x***^*′*^ in terms of Euclidean distance.

#### 2.2.2 NC-inspired representation learning

Although NC is an invariant phenomenon commonly observed in well-trained neural networks across various model architectures and classification tasks,^43^ it has been shown that this behavior only holds for class-balanced data.^53, 54^ In imbalanced, long-tail datasets, class center embeddings no longer naturally form an ETF, with embeddings of minor classes collapsing into similar directions.^53^ Since NC characterizes the structure of models with strong learning capability (i.e., near-zero training error) and generalizability,^43^ recent studies have explored various strategies to “force” a classifier to exhibit NC on imbalanced datasets, including loss regularization, gradient balancing, or optimal transport, which have been shown to induce NC and improve imbalanced classification performance, particularly in image classification.^53, 55–59^

Motivated by these findings, we aim to induce NC in imbalanced protein function annotation tasks. MSRep leverages ESM-1b,^45^ a pre-trained protein language model, whose sequence embeddings have been shown to be informative for protein function prediction.^28, 56^ MSRep refines ESM embeddings by re-projecting them into a new space that conforms to the *NC*_1_ and *NC*_2_ properties related to embedding learning, achieved by optimizing a loss function designed to shape the embedding space accordingly.

Specifically, given an input protein sequence ***s***, MSRep first applies ESM to generate an embedding ***x***^ESM^ with dimensionality *d*_ESM_ = 1280. Next, a trainable projection network *g*_*ω*_ transforms ***x***^ESM^ into a new embedding ***x*** in a space of dimensionality *d*. The projection network consists of fully connected layers (i.e., multi-layer perceptron, MLP), each followed by a LayerNorm, ReLU activation, and Dropout. To ensure NC properties (particularly *NC*_2_), the dimensionality *d* of the new embedding space must satisfy *d* ≥ *K* − 1.^43^ To induce NC in this *d*-dimensional space, we introduce two loss functions, inspired by the *NC*_1_ and *NC*_2_ properties, to constrain the learning of the sequence embeddings ***x***’s and a set of trainable embeddings 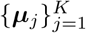, which act as the class centers. These loss functions are defined as:

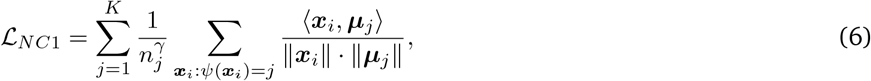

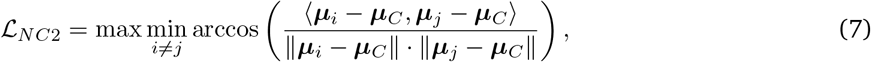

where *n*_*j*_ is the number of training samples in class *j, ψ*(·) maps a sequence to its true function label, *γ >* 0 is a hyperparameter controlling the weight of each class in 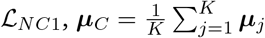 is the global mean of class centers, which is updated in each gradient step when calculating *ℒ*_*NC*2_.

Intuitively, *ℒ*_*NC*2_ ensures the *NC*_2_ property, encouraging 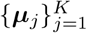 to behave like class centers in NC, forming an EFT after zero-centering. This is achieved by maximizing the minimum pairwise angle between any two class centers, ensuring they are maximally separated with equal-sized angles. *ℒ* _*NC*1_ enforces the *NC*_1_ property by pushing each sequence embedding ***x***_*i*_ toward its corresponding class center ***µ***_*j*_. The sum of cosine distances between each sequence embedding and its class center is calculated, weighted by the inverse class sizes, with the hyperparameter *γ* preventing larger classes from dominating *ℒ* _*NC*1_. Through hyperparameter tuning, we found that setting *γ* to 0.5 or 1.0 was robust across several protein function annotation tasks.

The final loss function is a linear combination of *ℒ*_*NC*1_ and *ℒ*_*NC*2_, controlled by a interpolation factor *β*:

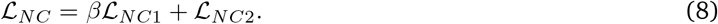

Through grid search on the validation set, *β* = 1.0 was found to be robust across all experiments. During training, only the MLP projection network *g*_*ω*_ and the trainable embeddings 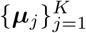 are optimized, while the ESM parameters are kept fixed.

By optimizing *ℒ*_*NC*_, MSRep learns an embedding space where all classes occupy equal portions of the space and are maximally separated (Fig. 1c). This regularized space enables tail-class proteins to be effectively classified, contrasting with conventional classification methods, where the vast majority of the leaned space is used to represent head-class proteins, whereas tail-class proteins are overwhelmed and occupy a small portion of the space (Fig. 1c). MSRep generalizes recent contrastive learning approaches^23, 28^ by optimizing group-wise distances instead of pairwise distances to organize the embedding space. Additionally, unlike many computer vision methods for imbalanced classification,^53, 55–59^ which incorporated *ℒ*_*NC*_ as a regularizer for cross-entropy loss, MSRep optimizes only *ℒ*_*NC*_ to mitigate the highly unstable interplay between these two losses during training for improved performance (see discussion in Supplementary Note A.5).

#### 2.2.3 Training and inference details

The original NC statement (Sec. 2.2.1) holds for *single-label, multi-class* classification tasks where each sample is assigned only one label from a total of *K* classes. However, protein function prediction is inherently a *multi-label, multi-class* classification task, where a protein can perform multiple functions and thus have more than one label. We used an approach that effectively handled the multi-label nature of the problem in our experiments: for a sample (***s***_*i*_, ***y***_*i*_) where the sequence ***s*** has multiple function labels 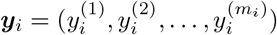, we replace it with multiple single-labeled samples 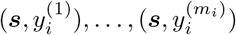.

For each function annotation task, we randomly withheld 10% of the training data as a validation set. MSRep was trained using the Adam optimizer^60^ and a learning rate of 1 *×* 10^−4^ for up to 10, 000 epochs. Early stopping was employed if the validation loss did not drop for 20 consecutive epochs. Task-specific hyperparameters were tuned on the respective validation set, and the final values used in our experiments are listed in Supplementary Table S3.

At prediction time, MSRep employed a nearest neighbor search approach, which has been shown effective in previous work.^23, 28^ Given a query protein, MSRep searches the training set for the protein with the smallest cosine distance to the query in the learned embedding space. The function annotations of the nearest neighbor were then transferred to the query protein as its predicted function labels.

To enhance prediction performance, we applied an ensemble scheme based on majority voting. Specifically, for each function annotation task, we trained five instances of MSRep, each initialized with a different random seed but with identical network architecture. At prediction time, each model predicted function labels for the query protein using the nearest-neighbor strategy. If a label was predicted by more than half of the individual models, it was included in the final predicted labels for the query protein. We observed that the MSRep’s ensemble model consistently outperformed its individual model (Supplementary Table S1).

## 3 Results

To evaluate the performance of MSRep, we constructed four benchmark datasets for different function annotation tasks: EC numbers, Gene3D codes, Pfam families, and GO terms (Fig. 1b). For each task, we compared MSRep against respective state-of-the-art methods (Supplementary Notes A.4).

### 3.1 Benchmark datasets

We utilized protein sequence and function annotation data in the Swiss-Prot database^61^ and implemented a time-based train-test split with a cutoff of May 25, 2022, simulating a realistic scenario by assessing the model’s generalizability on unseen proteins. Additionally, the test set was filtered by MMseqs^62^ to remove sequences with ≥ 50% sequence identity to any training sequence, resulting in test sets for the four tasks, denoted as EC, Gene3D, Pfam, GO-S50. For EC number prediction, we further collected the Price test set, a recognized challenging dataset derived from an experimental study,^63^ which includes sequences known to be incorrectly annotated in other databases.^49^ Detailed information on dataset construction and statistics is provided in Supplementary Notes A.1 and Supplementary Table S2.

### 3.2 MSRep improves protein function annotations

#### 3.2.1 EC numbers: Enzyme function prediction

The EC number ontology, which annotates enzyme functions, exhibits a highly imbalanced distribution. In our training set, among more than 5,000 unique EC numbers, fewer than 10% (509 EC numbers) are associated with over 84% proteins, while the remaining EC numbers have fewer than 100 associated proteins each (Fig. 1a). We trained MSRep on Swiss-Prot proteins annotated with EC numbers before 2022/05/25, and compared its performance with CLEAN,^28^ the current state-of-the-art enzyme function predictor, as well as four additional competitive baselines: Protein/Aspect-Vec,^23^ ProteInfer,^21^ and DeepEC.^27^

On the sequence identity-controlled test set EC-S50, MSRep significantly outperformed other methods across precision, recall, and F1 metrics (Fig 2a). To assess its performance on tail-class functions, we restricted the evaluation to function classes represented by fewer than 30 proteins in the training data. While most baselines showed a performance decrease, MSRep’s performance dropped less severely, maintaining a clear improvement in prediction accuracy (Fig 2b). We further delineated the analysis by evaluating methods in four bins of function classes, categorized by their occurrence (number of associated proteins) in the training set. MSRep showed consistent F1 improvements across all bins (Fig 2c). Remarkably, even for extremely underrepresented classes with fewer than 10 associated proteins, MSRep achieved a competitive F1 score exceeding 0.4, outperforming all other methods. We also benchmarked MSRep on the challenging Price-145 test set and found that it consistently outperformed existing methods on both the full test set (Fig S1a) and a subset of tail-class functions (Fig S1b).

**Figure 2:**
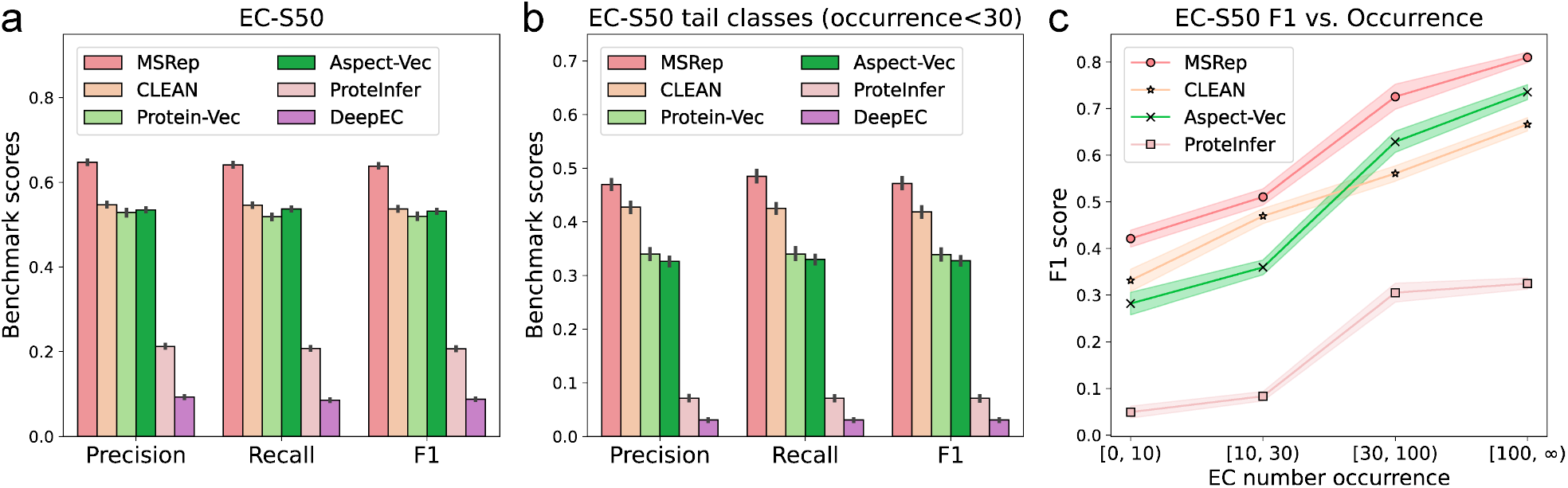
Performance evaluation on EC number prediction. (**a**) Precision, recall, and F1 scores (averaged across proteins) for each method evaluated on the full set of EC numbers in EC-S50. (**b**) Performance metrics for tail-class EC numbers with occurrence *<*30, where the occurrence is defined as the number of proteins associated with an EC number. (**c**) F1 scores across classes binned by their occurrence in the training data. Bar plots in **a-b** and the curves in **c** represented the mean*±*s.d. over ten sets of 90% bootstrapped proteins from EC-S50.

Since each EC number follows a four-digit hierarchical structure (e.g., 5.5.1.4), baselines like ProteInfer and DeepEC predict up to the 3rd level (i.e., 5.5.1.X) if they lack confidence in the 4th-level prediction. To ensure a fair comparison, we evaluated all methods based on exact-match accuracy at both the 3rd and 4th levels (Supplementary Notes A.2). Results showed that MSRep achieved more correct predictions at both levels, with fewer incorrect predictions compared to the baselines (Fig. S1c).

#### 3.2.2 Gene3D: Structure domain prediction

Gene3D^47^ is a database that provides structural domain family annotations for protein sequences based on the CATH ontology, which organizes protein structural domains into a four-level hierarchy: Class (C), Architecture (A), Topology/fold (T), and Homologous superfamily (H). Since proteins can harbor multiple structural domains, the Gene3D task represents a multi-label, multi-classification task over ~5, 000 classes.

We trained a MSRep model to predict Gene3D codes for protein sequences. On the Gene3D-S50 test set, MSRep outperformed competitive baselines, including Protein/Aspect-Vec and the established BLAST algorithm (Fig. 3a). The performance improvement was particularly notable when focusing on minority Gene3D classes with fewer than 30 occurrences in the training data, where MSRep achieved higher precision, recall, and F1 scores across all evaluated metrics (Fig. 3a).

**Figure 3:**
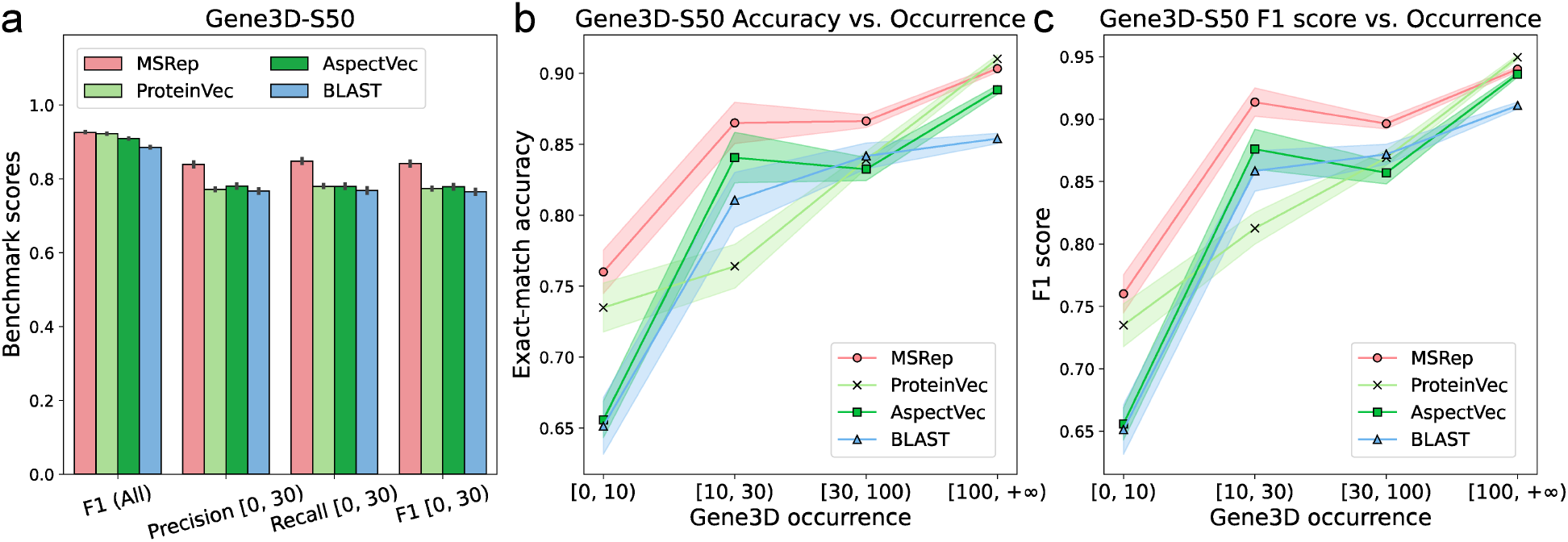
Performance evaluation on Gene3D prediction. (**a**) Precision, recall, and F1 scores for each method, evaluated on both the full set of classes and the tail classes (occurrence *<* 30) in Gene3D-S50. (**b-c**) Exact-match accuracy and F1 scores across classes binned by their occurrences in the training data. Bar plots in **a** and the curves in **b-c** represented the mean*±*s.d. over ten sets of 90% bootstrapped proteins from Gene3D-S50.

To further examine MSRep’s imbalanced learning ability, we evaluated its performance across different occurrence bins. For Gene3D codes with fewer than 100 associated proteins, MSRep achieved substantially higher exact-match accuracy and F1 scores than the baselines, demonstrating its robustness in predicting rare structural domain annotations. For frequent Gene3D codes with 100 or more occurrences, MSRep performed competitively, matching the accuracy of the strongest baseline methods. Note that the exact-match accuracy metric (defined in Supplementary A.2) only considers predictions correct if the entire set of labels matches the ground truth in both size and composition, which suggests the good performance of MSRep for multi-label classification.

#### 3.2.3 Pfam: Remote homology detection

Next, we trained MSRep to predict protein family annotations defined by the Pfam database.^35^ Our training set derived from Swiss-Prot includes 14,723 unique family labels, exhibiting a highly long-tail distribution, with nearly 90% of Pfam labels associated with fewer than 100 sequences (Fig. 1a). Since Pfam family labels are defined on the protein domain level, some previous works^31^ formulated this task as single-label classification using the subsequence in a protein corresponding to the domain as input. In contrast, we extend this formulation to a more general, multi-label classification setting, using full-length protein sequences to predict the Pfam labels for all domains presented in a protein. We compared MSRep to several strong baselines, including Protein/Aspect-Vec,^23^ which have shown state-of-the-art prediction performance, and ProtCNN/ProtENN,^31^ which are used to generate ML-predicted family labels in Pfam’s official releases.^36^

On the Pfam-S50 test set, MSRep consistently outperformed all baseline methods across classes with varying occurrence levels, achieving higher exact-match accuracy and F1 scores (Fig. 4a-b). Remote homology detection is particularly important in function annotation, as many proteins cannot be annotated solely through sequence alignment due to low sequence similarity to existing annotated proteins. To assess MSRep’s performance in this context, we further restricted the test set to sequences with *<* 30% sequence identity to the training data. While the F1 score of baselines like ProtENN and Aspect-Vec dropped considerably for these low-homology proteins, MSRep maintained significantly higher F1 scores, demonstrating resilience to low sequence identity (Fig. 4c). Moreover, when we varied the maximum sequence identity threshold, MSRep consistently outperformed the baselines across all cutoff values (Fig. 4c).

**Figure 4:**
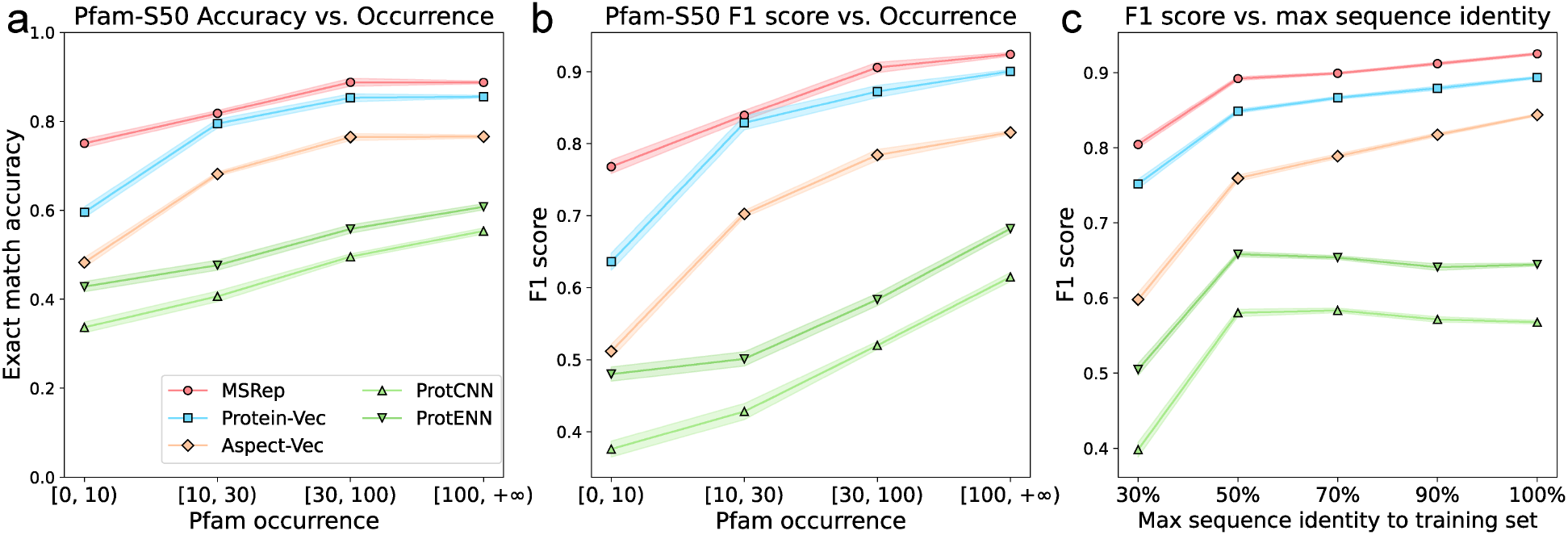
Performance evaluation on Pfam prediction. (**a-b**) Exact-match accuracy and F1 performance on the Pfam-S50 test set across different occurrence bins. (**c**) F1 scores across test sets with varying levels of maximum sequence identity to the training sequences. Line plots in this figure represent mean ± s.d. over ten sets of 90% bootstrapped samples from the test data.

Considering the number of unique Pfam labels is substantially larger than for EC numbers or Gene3D codes (approximately 15,000 v.s. 5,000; Supplementary Table S2), these results suggested that MSRep is scalable to large function annotation tasks and provides accurate predictions across diverse protein families.

#### 3.2.4 Gene Ontology: GO-term function annotation

To further evaluate the effectiveness of MSRep, we extended our benchmark to Gene Ontology (GO) annotations–a widely used ontology that describes gene and protein functions at multiple levels of granularity. Unlike EC numbers and Gene3D, which have well-defined four-level tree structures, the GO hierarchy is more complex due to variable depth for leaf nodes and that child terms may have multiple parent terms at various levels. Predicting GO terms for proteins is a longstanding challenge in computational biology and the focus of the community-wide Critical Assessment of Functional Annotation (CAFA).^33^

Following established practices in recent studies,^18, 22, 24^ we trained a separate MSRep model for each of the three main GO sub-ontologies: biological process (BP), molecular function (MF), and cellular component (CC). Each sub-ontology contains thousands of GO terms (ranges from 3,000 to 17,000), with a pronounced long-tail occurrence distribution (Fig. 1a). We compared MSRep against three recent high-performance GO predictors–AnnoPro, NetGO3.0, and Protein-Vec^23^–following the CAFA challenge’s setting^33, 64^ of evaluation metrics (Fmax and AUPR) and post-inference processing (Supplementary Notes A.3).

The results demonstrated MSRep’s strong performance in predicting GO terms (Table 1). Across 18 evaluations–covering 3 sub-ontologies, 3 class occurrence bins, and 2 metrics–MSRep achieved the highest scores in 16 cases and remained competitive in the remaining two. Its performance was especially pronounced in the tail-class regime, where it outperformed the second-best methods by 6.6%-28.3% in Fmax and 18.5%-92.6% in AUPR, indicating its ability to accurately predict GO annotations for understudied classes.

**Table 1:** Performance evaluation on GO term prediction. Fmax and AUPR scores for methods evaluated on the major three GO sub-ontologies: biological process (BP), molecular function (MF), and cellular component (CC). For each sub-ontology, the best performance is shown in bold, and the second best is underlined. Results are reported as mean*±*s.d. based on ten sets of 90% bootstrapped proteins from GO-S50. Abbreviations: Ont=Ontology.

| Ont | Method | [0, 30) |  | [30, 100) |  | [100, +∞) |  |
| --- | --- | --- | --- | --- | --- | --- | --- |
|  |  | Fmax | AUPR | Fmax | AUPR | Fmax | AUPR |
| BP | NetGO3.0 | 0.1209±.0037 | 0.0475±.0031 | 0.1809±.0066 | 0.0949±.0044 | 0.4960±.0033 | 0.4629±.0038 |
|  | AnnoPro | 0.0001±.0000 | 0.0000±.0000 | 0.0101±.0012 | 0.0005±.0001 | 0.3494±.0050 | 0.2922±.0039 |
|  | Protein-Vec | 0.1929±.0139 | 0.1189±.0113 | 0.2228±.0107 | 0.1325±.0075 | 0.5332±.0036 | 0.4389±.0037 |
|  | MSRep | **0.2056±.0125** | **0.1409±.0144** | **0.2949±.0152** | **0.1710±.0114** | **0.5731±.0028** | **0.5095±.0027** |
| MF | NetGO3.0 | 0.3169±.0131 | 0.1348±.0106 | **0.4555±.0118** | **0.3852±.0118** | 0.6286±.0022 | 0.6714±.0024 |
|  | AnnoPro | 0.0003±.0001 | 0.0001±.0000 | 0.1038±.0089 | 0.0242±.0063 | 0.6088±.0021 | 0.6176±.0026 |
|  | Protein-Vec | 0.2583±.0108 | 0.1203±.0063 | 0.3727±.0085 | 0.2939±.0074 | 0.7894±.0029 | 0.7798±.0034 |
|  | MSRep | **0.3395±.0079** | **0.2596±.0130** | 0.4094±.0131 | 0.3386±.0125 | **0.8152±.0027** | **0.8290±.0044** |
| CC | NetGO3.0 | 0.1455±.0106 | 0.0435±.0051 | 0.3786±.0079 | 0.1762±.0064 | 0.5708±.0022 | 0.5888±.0023 |
|  | AnnoPro | 0.0001±.0000 | 0.0000±.0000 | 0.0666±.0065 | 0.0269±.0047 | 0.5562±.0016 | 0.5160±.0020 |
|  | Protein-Vec | 0.2123±.0184 | 0.1272±.0165 | 0.3093±.0101 | 0.2587±.0165 | 0.6675±.0020 | 0.5641±.0039 |
|  | MSRep | **0.2724±.0224** | **0.2049±.0250** | **0.4322±.0126** | **0.3527±.0140** | **0.7210±.0022** | **0.7548±.0029** |

##### Performance evaluation summary

Overall, our experiments highlight the excellent predictive performance of MSRep across four diverse function annotation tasks. MSRep consistently outperformed state-of-the-are ML methods. Notably, it demonstrated robustness in predicting underrepresented function classes, an area where other methods experienced significant performance declines.

### 3.3 Analysis of MSRep’s learned representation space

Having established MSRep’s accuracy and generalizability for protein function annotation, we sought to understand the underlying reasons for its performance improvements, especially for minority function classes. The core idea of MSRep is to learn a regularized representation that conforms to the NC structure, ensuring that each function class, irrespective of its popularity in the training data, is modeled with equal representation capacity for accurate prediction. We hypothesize that this structure explains MSRep’s substantial performance gains for tail-class functions, as the representations for different functions are maximally separated, making them more discriminative than embeddings learned by conventional cross-entropy or contrastive learning losses.

To gain deeper insights into MSRep’s learned representation space, we conducted quantitative and visualization analyses, comparing the representation spaces learned by MSRep and a strong baseline, Aspect-Vec, on the EC number task. We first computed the intra-class and inter-class Euclidean embedding distances between protein pairs. MSRep more effectively pulled together proteins with the same function while pushing apart proteins with different functions, resulting in greater separation between intra- and inter-class distance distributions than observed for Aspect-Vec (Fig. 5a-b).

**Figure 5:**
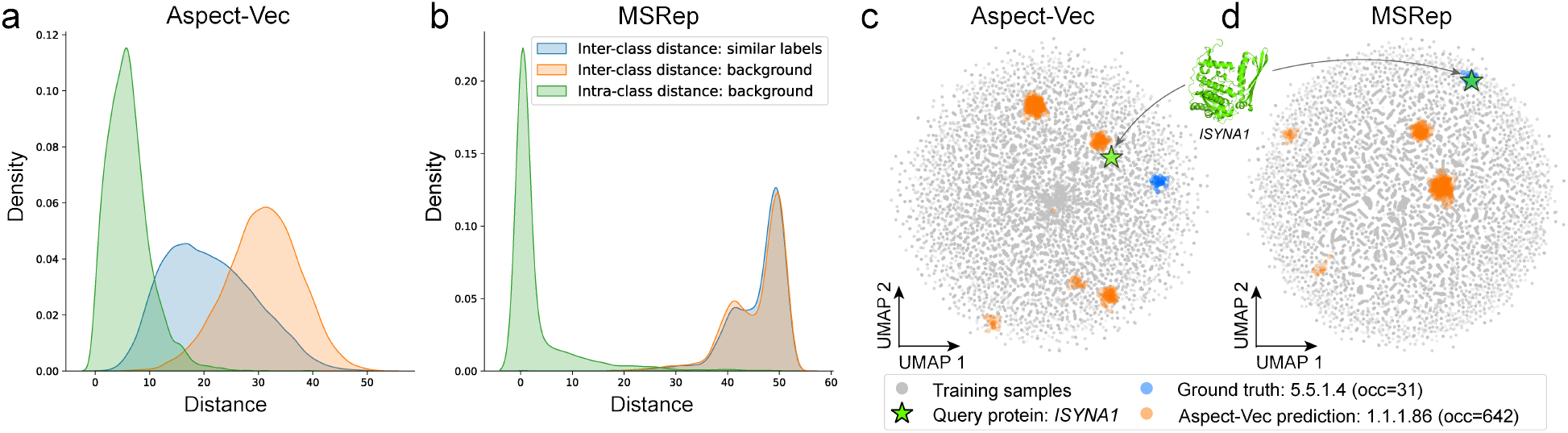
Representation learning of MSRep for EC number predictions. (**a-b**) Distribution of inter-class and intra-class Euclidean embedding distances for MSRep and Aspect-Vec. The “similar labels” category refers to pairs of EC numbers with the same first three digits but different fourth digits (e.g., 5.5.1.4 vs. 5.5.1.7). Background distributions show the pairwise distances of all intra-class pairs and 10^6^ randomly sampled inter-class pairs. (**c-d**) UMAP visualization of the representation spaces for Aspect-Vec and MSRep. Shown are the query protein *ISYNA1*, the training samples of its ground truth class (EC 5.5.1.4, occurrence = 31), and the training samples of the misclassified class predicted by Aspect-Vec (EC 1.1.1.86, occurrence = 642).

We further investigated pairwise embedding distance among proteins with highly similar enzymatic functions, defined as those sharing the same first three digits in their EC numbers but differing in the fourth digit (e.g., 5.5.1.4 vs. 5.5.1.7). Since these proteins are functionally similar, their sequences tend to be similar as well, which caused Aspect-Vec’s embeddings for these proteins to converge, shifting their pairwise distance distribution from the inter-class background toward the intra-class range Fig. 5a). This convergence made Aspect-Vec’s embeddings indistinguishable from other classes. In contrast, MSRep maintained separated embeddings for these functionally similar proteins, with their pairwise distance distribution almost identical to inter-class distance background (Fig. 5b), highlighting the effectiveness of MSRep’s NC-inspired loss in learning discriminative embeddings to inform function classification.

MSRep’s maximally spanning representations also translated to its performance gains, particularly in the long-tail regime where more discriminative embeddings facilitate classification. To illustrate this, we identified Inositol-3-phosphate synthase 1 (*ISYNA1*; UniProt ID: Q8A7J8), an enzyme with the EC number 5.5.1.4, which is an under-represented class with only 31 occurrences in the training data. While Aspect-Vec misclassified this protein to EC number 1.1.1.86, a well-represented class with 642 occurrences, MSRep classified it correctly. UMAP visualization of the embedding spaces learned by both methods revealed that MSRep’s NC-structured space correctly positioned this query protein within the cluster of proteins sharing the same EC number. In contrast, in Aspect-Vec’s space, the query protein was “attracted” to a large cluster of a head-class function, far from its functionally similar proteins.

Overall, these results suggested that the maximally spanning representation learned by MSRep produces more discriminative sequence embeddings, which are particularly informative for imbalanced classification.

## 4 Conclusion

We have presented MSRep, a deep learning framework for protein function annotation that addresses the challenge of data imbalance in protein function data. Systematic evaluations across four function annotation benchmarks showed that MSRep consistently outperformed state-of-the-art methods, particularly for under-represented function classes. While NC’s requirement for a high-dimensional embedding space (*d* ≥ *K* − 1) could lead to increased computational costs for tasks with numerous classes, this could potentially be mitigated by reducing the dimensionality, leveraging recent findings on generalized neural collapse in lower dimensions.^65^ Overall, MSRep provides a promising tool for sequence-based protein function annotation and can be adapted to various classification tasks in protein function studies.

## Acknowledgements

This work is supported in part by the National Institute of General Medical Sciences of the National Institutes of Health under award R35GM150890 and the GenAI Seed Grant Program from GaTech IDEaS. This work used the Delta GPU Supercomputer at NCSA of UIUC through allocation CIS230097 from the Advanced Cyberinfrastructure Coordination Ecosystem: Services & Support (ACCESS) program, which is supported by NSF grants #2138259, #2138286, #2138307, #2137603, and #2138296. The authors acknowledge the computational resources provided by Microsoft Azure through the Cloud Hub program at GaTech IDEaS and the Microsoft Accelerate Foundation Models Research (AFMR) program.

## A Supplementary Notes

### A.1 Benchmark datasets

We utilized the Swiss-Prot database,^61^ the expert-curated subset of UniProtKB,^51^ to retrieve protein sequences and their function annotations. We focused on four widely used annotation schemes: EC numbers, Gene3D codes, Pfam families, and GO terms. The original Swiss-Prot database contains approximately 570k expert-reviewed protein sequences. We excluded proteins whose sequence length is shorter than 10 amino acids or longer than 1022, as ESM-1b has an input sequence limit of 1,022 amino acids. This resulted in a filtered dataset of 551,965 sequences.

For each function annotation scheme, we constructed a benchmark dataset by retaining sequences labeled with that type of annotation. All tasks followed a time-based train-test split, using sequences deposited to Swiss-Prot by May 25, 2022, as the training set and sequences added after that date as the test set. We chose this cutoff date following the study of baseline methods used in this work,^23^ ensuring a fair comparison. Additionally, for each task’s test set, we applied MMseqs2^62^ to remove test sequences with ≥ 50% sequence identity to any training sequence, forming more challenging test sets labeled EC-S50, Gene3D-S50, Pfam-S50, and GO-S50, respectively. The statistics of the filtered time-split datasets are provided in Supplementary Table S2.

For EC numbers, we also included an independent test set, ‘Price’, derived from a previous experimental study.^63^ This dataset contains bacterial proteins for which automated annotation methods in databases like KEGG^49^ or SEED^66^ had assigned incorrect or inconsistent EC numbers. The Price dataset has been used as a challenging test set in prior works, including ProteInfer^21^ and CLEAN.^28^ From the original 149 sequences in Price, we excluded 4 sequences with EC numbers not present in our training set, resulting in a final test set of 145 proteins, denoted as Price-145.

### A.2 Evaluation metrics

The protein function annotation tasks we evaluated are all multi-label, multi-class classification tasks, where each sample may be associated with multiple labels. Below, we describe the metrics used in our evaluation:

#### Precision

For a single test sample, precision is defined as TP / (TP + FP), where TP is the number of true positives and FP is the number of false positives. Let *n*_test_ be the number of test samples and *n*_class_ be the number of total classes. We define two binary matrices, the ground truth matrix ***Y***_true_ and the prediction matrix ***Y***_pred_, both of size (*n*_test_, *n*_class_). The overall precision on the test set is calculated as the average precision across all test samples. Specifically, we utilized the <monospace>sklearn.metrics.precision_score </monospace>function from the <monospace>scikit-learn </monospace>package, with the parameter <monospace>average=‘samples’</monospace>. For precision on a subset of classes (e.g., within a certain training set occurrence range), we extracted the relevant columns of ***Y***_true_ and ***Y***_pred_ to form new matrices 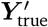 and 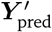, then computed precision in the same manner.

#### Recall

For a single test sample, recall is defined as TP / (TP + FN), where TP is the number of true positives and FN is the number of false negatives. Using ***Y***_true_ and ***Y***_pred_, we calculated the recall on the test set by averaging across all test samples. We used the <monospace>sklearn.metrics.recall_score </monospace>function in <monospace>scikit-learn </monospace>with <monospace>average=‘samples’</monospace>. The recall calculation on a subset of classes follows the same procedure as precision.

#### F1 score

For a single test sample, the F1 score is defined as 1 / (1 / precision + 1 / recall) = 2TP / (2TP + FP + FN), where TP, FP, and FN represent the counts of true positives, false positives, and false negatives, respectively. We calculated the F1 score using the same method as precision and recall, utilizing the <monospace>sklearn.metrics.f1_score </monospace>with <monospace>average=‘samples’</monospace>. F1 scores on a subset of classes were calculated similarly.

#### Exact-match accuracy

For multi-label classification, exact-match accuracy is defined as the percentage of test samples for which the predicted label set matches the ground-truth label sets exactly. For example, the *i*-th test sample is counted as correct if the *i*-th row of ***Y***_true_ and ***Y***_pred_ are identical. When evaluating exact-match accuracy on a subset of classes, we extracted the relevant columns from ***Y***_true_ and ***Y***_pred_ to form 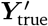 and 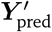, and computed accuracy on the new matrices.

#### Fmax score

Fmax score is the major evaluation metric used in the CAFA challenge,^33^ defined as the maximum F1 score obtained over a range of prediction thresholds. Let ***Y***_prob_ be the probability matrix of size (*n*_test_, *n*_class_), and ***Y***_true_ be the ground truth matrix. The *i*-th row of ***Y***_prob_ is the predicted probabilities for all classes for the *i*-th test sample. Given a threshold *ρ* ∈ [0, 1], we create a binary prediction matrix ***Y***_pred_ by setting entries in ***Y***_prob_ below *ρ* to 0 and the rest to 1. Using ***Y***_pred_ and ***Y***_true_, we calculate the F1 score at each threshold *ρ* using <monospace>sklearn.metrics.f1_score </monospace>with <monospace>average=‘micro’</monospace>. We use the micro average here to be consistent with the computation of AUPR, which uses the micro-averaged precision and recall. In our experiments, we used 101 thresholds from 0 to 1 with an increment of 0.01. For a subset of classes, the calculation follows the same procedure with subset matrices 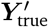 and 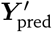.

#### AUPR

The area under the precision-recall curve (AUPR) is calculated by flattening ***Y***_prob_ and ***Y***_true_ into 1D arrays and computing the precision-recall curve using <monospace>sklearn.metrics.precision_recall_curve </monospace>and <monospace>sklearn.metrics.auc </monospace>in <monospace>scikit-learn</monospace>. For class subsets, we create new matrices 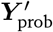 and 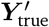 by selecting relevant columns, flattening them, and computing AUPR.

### A.3 Gene Ontology predictions

The evaluation scheme of the CAFA challenge,^33, 64^ which uses Fmax and AUPR as metrics, requires each method to associate a confidence score to each of its GO term predictions. For MSRep, this confidence score is calculated as the cosine similarity between the test protein embedding and the nearest reference embedding in the training set. Additionally, for each test protein and each GO term predicted by MSRep or baseline methods, we performed backpropagation to propagate the prediction to parent GO terms. Each parent GO term was assigned the same confidence score as the original predicted term, consistent with CAFA evaluation protocols.

### A.4 Baseline methods

#### Protein-Vec

Protein-Vec^23^ leverages protein language models and contrastive learning to generate multi-aspect protein representations. By pulling together embeddings of similar proteins and pushing apart those of distinct proteins, it captures multiple functional attributes (aspects), including EC numbers, Gene3D, Pfam, and Gene Ontology. For inference, Protein-Vec finds the nearest neighbor for each query sequence and transfers all annotations from the neighbor to the query. We used the pre-trained checkpoints of Protein-Vec, trained on Swiss-Prot sequences up to May 25, 2022, as a baseline for all tasks.

#### Aspect-Vec

Aspect-Vec,^23^ a single-aspect feature extractor model used in Protein-Vec’s construction, also employs contrastive learning but focuses on one function type at a time. We included Aspect-Vec as a baseline for EC numbers, Gene3D, and Pfam annotations.

#### BLAST

BLAST,^67^ a widely used sequence alignment tool, serves as a baseline for the Gene3D prediction task. Annotations for a query protein are transferred from the protein in the training set with the highest sequence similarity.

#### CLEAN

CLEAN^28^ is the state-of-the-art ML method for EC number prediction based on contrastive learning. For a fair comparison, we trained CLEAN on the same EC number training dataset as MSRep and performed inference using maximum separation as described in the original paper.

#### ProteInfer

ProteInfer^21^ is a deep convolutional neural network designed for EC number prediction. We used its pretrained checkpoints and the default threshold of 0.5, retaining only complete EC predictions for evaluation.

#### DeepEC

DeepEC^27^ utilizes three convolutional neural networks for EC number prediction. Using its pretrained model with the default threshold of 0.5, we retained all complete EC number predictions for evaluation.

#### ProtCNN and ProtENN

ProtCNN^31^ is a deep learning model for Pfam family predictions. It employs convolutional layers on one-hot encoded protein sequences to extract features and make Pfam predictions. ProtENN^31^ is an ensemble of 19 ProtCNN models, combining their outputs through majority voting. Notably, ProtCNN and ProtENN have been utilized to provide the ML-predicted family labels in the official Pfam releases.^36^

#### AnnoPro

AnnoPro^24^ is a deep learning algorithm for Gene Ontology (GO) prediction. It generates multi-scale protein representations, encodes them with convolutional and fully connected layers, and then decodes the features with a long short-term memory (LSTM) network for GO term predictions.

#### NetGO3.0

NetGO3.0^22^ is an automated function prediction framework for GO prediction. It integrates multiple prediction methods using GO label frequency, sequence alignment, k-mer features, protein domains, protein networks, and protein language model representations, combining them through a learning-to-rank algorithm.

### A.5 Methodological differences from previous methods

MSRep’s approach to learning a function-aware embedding space differs from recent contrastive learning methods that have achieved state-of-the-art protein function prediction.^23, 28^ Although seemly similar—both pulling functionally similar proteins together and pushing the functionally dissimilar proteins apart in the embedding space—contrastive learning relies on the optimization of many pairwise distances to organize the entire embedding space. In contrast, MSRep explicitly constructs a global “scaffolding backbone” of the embedding space, i.e., the ETF, and optimizes the sequence embeddings to conform to this structure.

Additionally, MSRep also differs from existing methods that induce NC to improve imbalanced classification in computer vision tasks.^53, 55–59^ These methods typically incorporate the NC loss *ℒ*_*NC*_ as a regularization term alongside the standard cross-entropy loss. However, the protein function annotation datasets (e.g., GO terms) considered in this work are far more diverse and imbalanced (~10^3^ − 10^4^ classes) than the image datasets (~10^1^ − 10^2^ classes) used in these studies. The dynamic interplay between the cross-entropy loss and the *ℒ*_*NC*_ loss is highly unstable during model optimization. As a result, MSRep optimizes only the *ℒ*_*NC*_ loss to learn the NC-structured embedding space and utilizes an effective nearest search scheme in this space for function annotation.

## B Supplementary Tables

**Table S1:** Comparison of individual versus ensemble models of MSRep. The metrics are evaluated on all classes within the task. Results are reported as mean ± s.d. over ten sets of 90% bootstrapped proteins from the corresponding test set. For each task, the entries with better performance are bolded. The accuracy listed in the table is the exact-match accuracy. Abbreviations: BP=biological process; MF=molecular function; CC=cellular component.

| Task | Method | Precision | Recall | F1 | Accuracy | Fmax | AUPR |
| --- | --- | --- | --- | --- | --- | --- | --- |
| EC (S50) | Individual | 0.6453±.0092 | 0.6344±.0100 | 0.6344±.0095 | **0.5888**±.0095 | - | - |
| | Ensemble | **0.6469**±.0081 | **0.6408**±.0090 | **0.6381**±.0084 | 0.5879±.0087 | - | - |
| EC (Price) | Individual | 0.3830±.0115 | **0.4592**±.0124 | 0.4046±.0115 | 0.3293±.0123 | - | - |
| | Ensemble | **0.3927**±.0116 | 0.4585±.0120 | **0.4105**±.0117 | **0.3515**±.0119 | - | - |
| Gene3D | Individual | 0.9181±.0022 | 0.9250±.0017 | 0.9151±.0016 | 0.8557±.0017 | - | - |
| | Ensemble | **0.9338**±.0024 | **0.9309**±.0017 | **0.9262**±.0020 | **0.8693**±.0017 | - | - |
| Pfam | Individual | 0.8953±.0029 | 0.9011±.0027 | 0.8898±.0027 | 0.8151±.0035 | - | - |
| | Ensemble | **0.8959**±.0027 | **0.9017**±.0028 | **0.8908**±.0027 | **0.8193**±.0036 | - | - |
| GO (BP) | Individual | - | - | - | - | 0.5438±.0025 | 0.4404±.0025 |
| | Ensemble | - | - | - | - | **0.5553**±.0033 | **0.4859**±.0033 |
| GO (MF) | Individual | - | - | - | - | 0.7596±.0027 | 0.7389±.0044 |
| | Ensemble | - | - | - | - | **0.8024**±.0027 | **0.8130**±.0043 |
| GO (CC) | Individual | - | - | - | - | 0.6692±.0022 | 0.6323±.0018 |
| | Ensemble | - | - | - | - | **0.7121**±.0021 | **0.7435**±.0030 |

**Table S2:** Statistics of the benchmarking datasets for the four protein function annotation tasks. For the three sub-ontologies BP, MF, and CC of Gene Ontology, the ‘Number of labels’ column indicates the count of leaf GO terms in the training set, i.e., those with no descendant terms.

| Task | Number of labels | Total data | Training data | Test data | Test data (S50) |
| --- | --- | --- | --- | --- | --- |
| EC number | 5,338 | 228,410 | 227,839 | 571 | 281 |
| Gene3D | 4,927 | 442,801 | 440,743 | 2,058 | 1,053 |
| Pfam | 14,723 | 523,041 | 520,636 | 2,405 | 1,207 |
| GO (BP) | 16,919 | 433,366 | 431,345 | 2,021 | 1,143 |
| GO (MF) | 7,900 | 468,451 | 466,029 | 2,422 | 1,353 |
| GO (CC) | 2,746 | 411,491 | 409,282 | 2,209 | 1,178 |

**Table S3:** Model architectures and hyperparameters for each function annotation task. The ‘num_layers’ column specifies the number of linear layers used in the projection network. ‘hidden_dims’ lists the hidden dimensions of the intermediate linear layers, and ‘out_dims’ represents the output dimensions of the projection network. All the hyperparameters were tuned and selected based on the performance on validation set.

| task | num_classes | num_layers | hidden_dims | out_dims | $\gamma$ | batch_size |
| --- | --- | --- | --- | --- | --- | --- |
| EC | 5,338 | 4 | [5120, 5120, 5120] | 5,400 | 0.5 | 10,000 |
| Gene3D | 4,927 | 3 | [5120, 5120] | 5,000 | 1.0 | 10,000 |
| Pfam | 14,723 | 2 | [5120] | 15,000 | 1.0 | 20,000 |
| GO (BP) | 16,919 | 2 | [2560] | 17,000 | 1.0 | 30,000 |
| GO (MF) | 7,900 | 3 | [5120, 5120] | 8,000 | 0.5 | 20,000 |
| GO (CC) | 2,746 | 3 | [5120, 5120] | 3,000 | 0.5 | 10,000 |

## C Supplementary Figures

**Figure S1:**
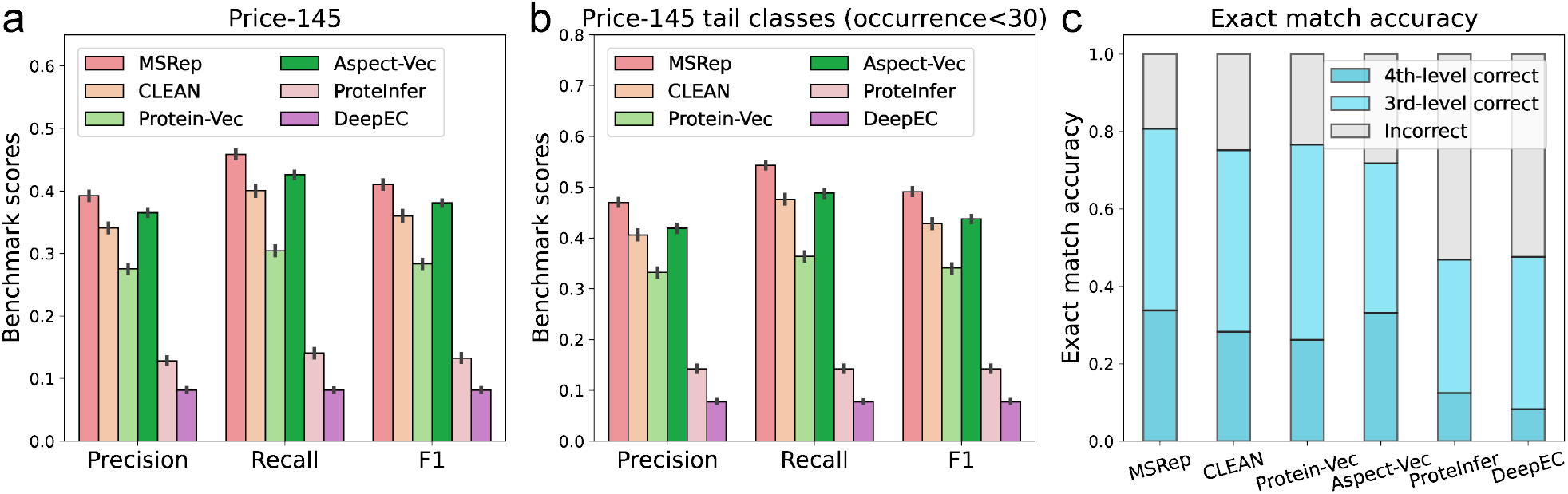
EC number prediction performance of MSRep on Price-145 test set. (**a**) Evaluation results across all EC classes. (**b**) Evaluation results on tail classes with training set occurrences below 30. (**c**) Exact-match accuracy results for the prediction of 4th-level and 3rd-level EC numbers. Bar plots in **a-b** represented the mean ± s.d. over ten sets of 90% bootstrapped proteins from the Price-145 test set.

## References

[1] Valerie Wood, Antonia Lock, Midori A Harris, Kim Rutherford, Jürg Bähler, and Stephen G Oliver. Hidden in plain sight: what remains to be discovered in the eukaryotic proteome? Open biology, 9(2):180241, 2019.

[2] Thomas Stoeger, Martin Gerlach, Richard I Morimoto, and Luís A Nunes Amaral. Large-scale investigation of the reasons why potentially important genes are ignored. PLoS biology, 16(9):e2006643, 2018.

[3] Winston A Haynes, Aurelie Tomczak, and Purvesh Khatri. Gene annotation bias impedes biomedical research. Scientific reports, 8(1):1362, 2018.

[4] The UniProt Consortium. Uniprot: the universal protein knowledgebase in 2023. Nucleic acids research, 51(D1):D523–D531, 2023.

[5] Yaan J Jang, Qi-Qi Qin, Si-Yu Huang, Arun T John Peter, Xue-Ming Ding, and Benoît Kornmann. Accurate prediction of protein function using statistics-informed graph networks. Nature Communications, 15(1):6601, 2024.

[6] Swati Sinha, Birgit Eisenhaber, Lars Juhl Jensen, Bharata Kalbuaji, and Frank Eisenhaber. Darkness in the human gene and protein function space: widely modest or absent illumination by the life science literature and the trend for fewer protein function discoveries since 2000. Proteomics, 18(21-22):1800093, 2018.

[7] Marian Breuer, Tyler M Earnest, Chuck Merryman, Kim S Wise, Lijie Sun, Michaela R Lynott, Clyde A Hutchison, Hamilton O Smith, John D Lapek, David J Gonzalez, et al. Essential metabolism for a minimal cell. Elife, 8:e36842, 2019.

[8] Tudor I Oprea, Cristian G Bologa, Søren Brunak, Allen Campbell, Gregory N Gan, Anna Gaulton, Shawn M Gomez, Rajarshi Guha, Anne Hersey, Jayme Holmes, et al. Unexplored therapeutic opportunities in the human genome. Nature reviews Drug discovery, 17(5):317–332, 2018.

[9] Ian Dunham. Human genes: Time to follow the roads less traveled? PLoS biology, 16(9):e3000034, 2018.

[10] Aled M Edwards, Ruth Isserlin, Gary D Bader, Stephen V Frye, Timothy M Willson, and Frank H Yu. Too many roads not taken. Nature, 470(7333):163–165, 2011.

[11] Georg Kustatscher, Tom Collins, Anne-Claude Gingras, Tiannan Guo, Henning Hermjakob, Trey Ideker, Kathryn S Lilley, Emma Lundberg, Edward M Marcotte, Markus Ralser, et al. An open invitation to the understudied proteins initiative. Nature Biotechnology, 40(6):815–817, 2022.

[12] Georg Kustatscher, Tom Collins, Anne-Claude Gingras, Tiannan Guo, Henning Hermjakob, Trey Ideker, Kathryn S Lilley, Emma Lundberg, Edward M Marcotte, Markus Ralser, et al. Understudied proteins: opportunities and challenges for functional proteomics. Nature Methods, 19(7):774–779, 2022.

[13] Stephen F Altschul, Thomas L Madden, Alejandro A Schäffer, Jinghui Zhang, Zheng Zhang, Webb Miller, and David J Lipman. Gapped blast and psi-blast: a new generation of protein database search programs. Nucleic acids research, 25(17):3389–3402, 1997.

[14] Johannes Söding. Protein homology detection by hmm–hmm comparison. Bioinformatics, 21(7):951– 960, 2005.

[15] Maxat Kulmanov, Mohammed Asif Khan, and Robert Hoehndorf. Deepgo: predicting protein functions from sequence and interactions using a deep ontology-aware classifier. Bioinformatics, 34(4):660–668, 2018.

[16] Renzhi Cao, Colton Freitas, Leong Chan, Miao Sun, Haiqing Jiang, and Zhangxin Chen. Prolango: protein function prediction using neural machine translation based on a recurrent neural network. Molecules, 22(10):1732, 2017.

[17] Balázs Szalkai and Vince Grolmusz. Near perfect protein multi-label classification with deep neural networks. Methods, 132:50–56, 2018.

[18] Maxat Kulmanov and Robert Hoehndorf. Deepgoplus: improved protein function prediction from sequence. Bioinformatics, 36(2):422–429, 2020.

[19] Nils Strodthoff, Patrick Wagner, Markus Wenzel, and Wojciech Samek. Udsmprot: universal deep sequence models for protein classification. Bioinformatics, 36(8):2401–2409, 2020.

[20] Vladimir Gligorijević, P Douglas Renfrew, Tomasz Kosciolek, Julia Koehler Leman, Daniel Berenberg, Tommi Vatanen, Chris Chandler, Bryn C Taylor, Ian M Fisk, Hera Vlamakis, et al. Structure-based protein function prediction using graph convolutional networks. Nature communications, 12(1):3168, 2021.

[21] Theo Sanderson, Maxwell L Bileschi, David Belanger, and Lucy J Colwell. Proteinfer, deep neural networks for protein functional inference. Elife, 12:e80942, 2023.

[22] Shaojun Wang, Ronghui You, Yunjia Liu, Yi Xiong, and Shanfeng Zhu. Netgo 3.0: protein language model improves large-scale functional annotations. Genomics, Proteomics & Bioinformatics, 21(2):349– 358, 2023.

[23] Tymor Hamamsy, Meet Barot, James T Morton, Martin Steinegger, Richard Bonneau, and Kyunghyun Cho. Learning sequence, structure, and function representations of proteins with language models. bioRxiv, 2023.

[24] Lingyan Zheng, Shuiyang Shi, Mingkun Lu, Pan Fang, Ziqi Pan, Hongning Zhang, Zhimeng Zhou, Hanyu Zhang, Minjie Mou, Shijie Huang, et al. Annopro: a strategy for protein function annotation based on multi-scale protein representation and a hybrid deep learning of dual-path encoding. Genome biology, 25(1):41, 2024.

[25] Yu Li, Sheng Wang, Ramzan Umarov, Bingqing Xie, Ming Fan, Lihua Li, and Xin Gao. Deepre: sequence-based enzyme ec number prediction by deep learning. Bioinformatics, 34(5):760–769, 2018.

[26] Zhenzhen Zou, Shuye Tian, Xin Gao, and Yu Li. mldeepre: multi-functional enzyme function prediction with hierarchical multi-label deep learning. Frontiers in genetics, 9:714, 2019.

[27] Jae Yong Ryu, Hyun Uk Kim, and Sang Yup Lee. Deep learning enables high-quality and high-throughput prediction of enzyme commission numbers. Proceedings of the National Academy of Sciences, 116(28):13996–14001, 2019.

[28] Tianhao Yu, Haiyang Cui, Jianan Canal Li, Yunan Luo, Guangde Jiang, and Huimin Zhao. Enzyme function prediction using contrastive learning. Science, 379(6639):1358–1363, 2023.

[29] Gi Bae Kim, Ji Yeon Kim, Jong An Lee, Charles J Norsigian, Bernhard O Palsson, and Sang Yup Lee. Functional annotation of enzyme-encoding genes using deep learning with transformer layers. Nature Communications, 14(1):7370, 2023.

[30] Yidong Song, Qianmu Yuan, Sheng Chen, Yuansong Zeng, Huiying Zhao, and Yuedong Yang. Accurately predicting enzyme functions through geometric graph learning on esmfold-predicted structures. Nature Communications, 15(1):8180, 2024.

[31] Maxwell L Bileschi, David Belanger, Drew H Bryant, Theo Sanderson, Brandon Carter, D Sculley, Alex Bateman, Mark A DePristo, and Lucy J Colwell. Using deep learning to annotate the protein universe. Nature Biotechnology, 40(6):932–937, 2022.

[32] Konstantin Schütze, Michael Heinzinger, Martin Steinegger, and Burkhard Rost. Nearest neighbor search on embeddings rapidly identifies distant protein relations. Frontiers in Bioinformatics, 2:1033775, 2022.

[33] Naihui Zhou, Yuxiang Jiang, Timothy R Bergquist, Alexandra J Lee, Balint Z Kacsoh, Alex W Crocker, Kimberley A Lewis, George Georghiou, Huy N Nguyen, Md Nafiz Hamid, et al. The cafa challenge reports improved protein function prediction and new functional annotations for hundreds of genes through experimental screens. Genome biology, 20:1–23, 2019.

[34] Huiying Yan, Shaojun Wang, Hancheng Liu, Hiroshi Mamitsuka, and Shanfeng Zhu. Goretriever: reranking protein-description-based go candidates by literature-driven deep information retrieval for protein function annotation. Bioinformatics, 40(Supplement_2):ii53–ii61, 2024.

[35] Alex Bateman, Lachlan Coin, Richard Durbin, Robert D Finn, Volker Hollich, Sam Griffiths-Jones, Ajay Khanna, Mhairi Marshall, Simon Moxon, Erik LL Sonnhammer, et al. The pfam protein families database. Nucleic acids research, 32(Suppl_1):D138–D141, 2004.

[36] Pfam. Google Research Team bring Deep Learning to Pfam — xfam.wordpress.com. https://xfam.wordpress.com/2021/03/24/google-research-team-bring-deep-learning-to-pfam/, 2021. [Accessed 17-Oct-2024].

[37] UniProt. Protein natural language model. https://www.uniprot.org/help/ProtNLM, 2022. [Accessed 17-Oct-2024].

[38] Maxat Kulmanov and Robert Hoehndorf. Deepgozero: improving protein function prediction from sequence and zero-shot learning based on ontology axioms. Bioinformatics, 38(Supplement_1):i238–i245, 2022.

[39] Rachael P Huntley, Tony Sawford, Prudence Mutowo-Meullenet, Aleksandra Shypitsyna, Carlos Bonilla, Maria J Martin, and Claire O’Donovan. The goa database: gene ontology annotation updates for 2015. Nucleic acids research, 43(D1):D1057–D1063, 2015.

[40] Barrett AJ. Nomenclature committee of the international union of biochemistry and molecular biology (nc-iubmb). enzyme nomenclature. recommendations 1992. Eur J Biochem, 1992.

[41] Lisa Van den Broeck, Dinesh Kiran Bhosale, Kuncheng Song, Cássio Flavio Fonseca de Lima, Michael Ashley, Tingting Zhu, Shanshuo Zhu, Brigitte Van De Cotte, Pia Neyt, Anna C Ortiz, et al. Functional annotation of proteins for signaling network inference in non-model species. Nature Communications, 14(1):4654, 2023.

[42] Jiaqi Luo and Yunan Luo. Contrastive learning of protein representations with graph neural networks for structural and functional annotations. In PACIFIC SYMPOSIUM ON BIOCOMPUTING 2023: Kohala Coast, Hawaii, USA, 3–7 January 2023, pages 109–120. World Scientific, 2022.

[43] Vardan Papyan, XY Han, and David L Donoho. Prevalence of neural collapse during the terminal phase of deep learning training. Proceedings of the National Academy of Sciences, 117(40):24652–24663, 2020.

[44] Vignesh Kothapalli. Neural collapse: A review on modelling principles and generalization. arXiv preprint 2206.04041, 2022.

[45] Alexander Rives, Joshua Meier, Tom Sercu, Siddharth Goyal, Zeming Lin, Jason Liu, Demi Guo, Myle Ott, C Lawrence Zitnick, Jerry Ma, et al. Biological structure and function emerge from scaling unsupervised learning to 250 million protein sequences. Proceedings of the National Academy of Sciences, 118(15):e2016239118, 2021.

[46] Gene Ontology Consortium. The gene ontology (go) database and informatics resource. Nucleic acids research, 32(Suppl_1):D258–D261, 2004.

[47] Tony E Lewis, Ian Sillitoe, Natalie Dawson, Su Datt Lam, Tristan Clarke, David Lee, Christine Orengo, and Jonathan Lees. Gene3d: extensive prediction of globular domains in proteins. Nucleic acids research, 46(D1):D435–D439, 2018.

[48] Damian Szklarczyk, Rebecca Kirsch, Mikaela Koutrouli, Katerina Nastou, Farrokh Mehryary, Radja Hachilif, Annika L Gable, Tao Fang, Nadezhda T Doncheva, Sampo Pyysalo, et al. The string database in 2023: protein–protein association networks and functional enrichment analyses for any sequenced genome of interest. Nucleic acids research, 51(D1):D638–D646, 2023.

[49] Hiroyuki Ogata, Susumu Goto, Wataru Fujibuchi, and Minoru Kanehisa. Computation with the kegg pathway database. Biosystems, 47(1-2):119–128, 1998.

[50] Michael Heinzinger, Maria Littmann, Ian Sillitoe, Nicola Bordin, Christine Orengo, and Burkhard Rost. Contrastive learning on protein embeddings enlightens midnight zone. NAR genomics and bioinformatics, 4(2):lqac043, 2022.

[51] Emmanuel Boutet, Damien Lieberherr, Michael Tognolli, Michel Schneider, Parit Bansal, Alan J Bridge, Sylvain Poux, Lydie Bougueleret, and Ioannis Xenarios. Uniprotkb/swiss-prot, the manually annotated section of the uniprot knowledgebase: how to use the entry view. Plant bioinformatics: methods and protocols, pages 23–54, 2016.

[52] Thomas Strohmer and Robert W Heath Jr. Grassmannian frames with applications to coding and communication. Applied and computational harmonic analysis, 14(3):257–275, 2003.

[53] Cong Fang, Hangfeng He, Qi Long, and Weijie J Su. Exploring deep neural networks via layer-peeled model: Minority collapse in imbalanced training. Proceedings of the National Academy of Sciences, 118(43):e2103091118, 2021.

[54] Christos Thrampoulidis, Ganesh Ramachandra Kini, Vala Vakilian, and Tina Behnia. Imbalance trouble: Revisiting neural-collapse geometry. Advances in Neural Information Processing Systems, 35:27225– 27238, 2022.

[55] Yibo Yang, Shixiang Chen, Xiangtai Li, Liang Xie, Zhouchen Lin, and Dacheng Tao. Inducing neural collapse in imbalanced learning: Do we really need a learnable classifier at the end of deep neural network? Advances in neural information processing systems, 35:37991–38002, 2022.

[56] Xuantong Liu, Jianfeng Zhang, Tianyang Hu, He Cao, Yuan Yao, and Lujia Pan. Inducing neural collapse in deep long-tailed learning. In International Conference on Artificial Intelligence and Statistics, pages 11534–11544. PMLR, 2023.

[57] Wanli Hong and Shuyang Ling. Neural collapse for unconstrained feature model under cross-entropy loss with imbalanced data. arXiv preprint 2309.09725, 2023.

[58] Gao Peifeng, Qianqian Xu, Peisong Wen, Zhiyong Yang, Huiyang Shao, and Qingming Huang. Feature directions matter: Long-tailed learning via rotated balanced representation. In International Conference on Machine Learning, pages 27542–27563. PMLR, 2023.

[59] Jintong Gao, He Zhao, Dan dan Guo, and Hongyuan Zha. Distribution alignment optimization through neural collapse for long-tailed classification. In Forty-first International Conference on Machine Learning, 2024.

[60] Diederik P Kingma. Adam: A method for stochastic optimization. arXiv preprint 1412.6980, 2014.

[61] Brigitte Boeckmann, Amos Bairoch, Rolf Apweiler, Marie-Claude Blatter, Anne Estreicher, Elisabeth Gasteiger, Maria J Martin, Karine Michoud, Claire O’Donovan, Isabelle Phan, et al. The swiss-prot protein knowledgebase and its supplement trembl in 2003. Nucleic acids research, 31(1):365–370, 2003.

[62] Martin Steinegger and Johannes Söding. Mmseqs2 enables sensitive protein sequence searching for the analysis of massive data sets. Nature biotechnology, 35(11):1026–1028, 2017.

[63] Morgan N Price, Kelly M Wetmore, R Jordan Waters, Mark Callaghan, Jayashree Ray, Hualan Liu, Jennifer V Kuehl, Ryan A Melnyk, Jacob S Lamson, Yumi Suh, et al. Mutant phenotypes for thousands of bacterial genes of unknown function. Nature, 557(7706):503–509, 2018.

[64] Damiano Piovesan, Davide Zago, Parnal Joshi, M Clara De Paolis Kaluza, Mahta Mehdiabadi, Rashika Ramola, Alexander Miguel Monzon, Walter Reade, Iddo Friedberg, Predrag Radivojac, et al. Cafa-evaluator: a python tool for benchmarking ontological classification methods. Bioinformatics Advances, 4(1):vbae043, 2024.

[65] Jiachen Jiang, Jinxin Zhou, Peng Wang, Qing Qu, Dustin Mixon, Chong You, and Zhihui Zhu. Generalized neural collapse for a large number of classes. arXiv preprint 2310.05351, 2023.

[66] Ross Overbeek, Tadhg Begley, Ralph M Butler, Jomuna V Choudhuri, Han-Yu Chuang, Matthew Cohoon, Valérie de Crécy-Lagard, Naryttza Diaz, Terry Disz, Robert Edwards, et al. The subsystems approach to genome annotation and its use in the project to annotate 1000 genomes. Nucleic acids research, 33(17):5691–5702, 2005.

[67] Stephen F Altschul, Warren Gish, Webb Miller, Eugene W Myers, and David J Lipman. Basic local alignment search tool. Journal of molecular biology, 215(3):403–410, 1990.

